# Mef2 induction of the immediate early gene *Hr38/Nr4a* is terminated by Sirt1 to promote ethanol tolerance

**DOI:** 10.1101/205666

**Authors:** Pratik Adhikari, Donnoban Orozco, Fred W. Wolf

## Abstract

Drug naïve animals given a single dose of ethanol show changed responses to subsequent doses, including the development of ethanol tolerance and ethanol preference. These simple forms of behavioral plasticity are due in part to changes in gene expression and neuronal properties. Surprisingly little is known about how ethanol initiates changes in gene expression or what the changes do. Here we demonstrate a role in ethanol plasticity for Hr38, the sole Drosophila homolog of the mammalian Nr4a1/2/3 class of immediate early response transcription factors. Acute ethanol exposure induces transient expression of *Hr38* and other immediate early neuronal activity genes. Ethanol activates the Mef2 transcriptional activator to induce *Hr38*, and the Sirt1 histone/protein deacetylase terminates *Hr38* induction. Loss of *Hr38* decreases ethanol tolerance and causes precocious but short-lasting ethanol preference. Similarly, reduced Mef2 activity in all neurons or specifically in the mushroom body α/β neurons decreases ethanol tolerance; Sirt1 promotes ethanol tolerance in these same neurons. Genetically decreasing *Hr38* expression levels in *Sirt1* null mutants restores ethanol tolerance, demonstrating that both induction and termination of *Hr38* expression are important for behavioral plasticity to proceed. These data demonstrate that Hr38 functions as an immediate early transcription factor that promotes ethanol behavioral plasticity.

## Introduction

Ethanol, one of the most widely used and frequently abused addictive drugs, is a small molecule that diffuses rapidly throughout the body and that binds to an as yet incompletely defined spectrum of molecules. Its effects vary based on dose, exposure time and pattern, an individuals’ history of intake, and their genetic makeup. This complexity of action has hampered progress in reducing the prevalence of alcohol use disorders through rational interventions (Edenberg & Foroud 2013).

One approach forward is to define, in detail, the stimulus-response relationship for ethanol in ethanol naïve animals. While the first ethanol exposure rarely leads directly to alcoholism, it does cause changes in behavior that reflect changes in brain function; these changes provide an altered substrate for subsequent intake and promote addiction risk. Furthermore, many genes are oppositely regulated by acute ethanol exposure and in ethanol withdrawal (Palmisano & Pandey 2017). Practically, this suggests that detailed mechanistic understanding of acute ethanol exposure action, especially when coupled to measures of behavioral plasticity, will provide insight into the more complex mechanisms underlying addiction. One form of ethanol-induced behavioral plasticity is tolerance, the acquired resistance to the inebriating and sedating properties of ethanol (Fadda & Rossetti 1998). Ethanol tolerance facilitates increased intake, a risk factor for later developing alcohol use disorders.

Drugs of abuse, including ethanol, cause changes in gene expression in the brain that can alter the properties of the brain. *Drosophila melanogaster* is a useful organism for defining how acute ethanol alters behavior through gene regulation (Kaun *et al.* 2012). In Drosophila, as in mammals, acute ethanol exposure progressively stimulates locomotion, motor incoordination, and sedation. Ethanol exposure also induces ethanol tolerance, ethanol preference, ethanol reward, and signs of ethanol withdrawal. Acute ethanol exposure causes marked changes in gene expression, and some ethanol-regulated genes have been shown to be critical for ethanol-induced behavioral plasticity (Morozova *et al.* 2007; Urizar *et al.* 2007; Ghezzi *et al.* 2010, 2013; Kong *et al.* 2010; Engel *et al.* 2016).

Neural activity and drugs of abuse induce immediate early genes in the nervous system, including transcription factors like Fos that are important for driving programs of gene expression (Sheng & Greenberg 1990; Robison & Nestler 2011). Neural activity in Drosophila also induces immediate early genes (Guan *et al.* 2005; Chen *et al.* 2016; Ghezzi *et al.* 2016). As yet, miRNAs are the only class of immediate early genes that have been studied in the context of ethanol behaviors in Drosophila (Ghezzi *et al.* 2016). Understanding how ethanol regulates immediate early response gene expression, and the consequences of their regulation, can help define how ethanol alters neural function and behavior.

Hr38 is a Drosophila immediate early response gene that is the sole homolog of the mammalian Nr4a1/Nr4a2/Nr4a3 gene family. Hr38 is strongly and consistently induced by artificial neural activation (Fujita *et al.* 2013; Chen *et al.* 2016). It functions in various processes in both development and adulthood, including ecdysis, carbohydrate storage, and circadian rhythms (Baker *et al.* 2003; Ruaud *et al.* 2011; Mezan *et al.* 2016). Here we find that *Hr38* and other immediate early genes are induced by ethanol, we define the mechanisms of *Hr38* induction, and we demonstrate that *Hr38* and its regulators function in the fly brain to promote the development of ethanol tolerance. These findings define early steps in gene regulation by ethanol that are important for the expression of ethanol-induced behavioral plasticity.

## Materials and Methods

### Drosophila Culturing and Strains

All strains were outcrossed for at least five generations to the Berlin genetic background carrying the *w1118* genetic marker mutation. The genetic background strain was used as an experimental control. Flies were cultured on standard cornmeal/molasses/yeast medium at 25°C and 70% relative humidity with a 12/12 hr light/dark schedule. Strains used in this study were *UAS-Mef2.EnR* from Justin Blau (Blanchard *et al.* 2010), *Hr38^y214^* from Carl Thummel (Kozlova *et al.* 2009), *MB-Gal80* from Scott Waddell, University of Oxford, Oxford, UK, *UAS-Mef2.IR* (Vienna Drosophila Resource Center, v15549, v15550), *UAS-Hr38.GFP* (Bloomington Drosophila Stock Center (BDSC) #38651), *17d-Gal4* (BDSC #51631), *elav-Gal4* (*c155*, BDSC #458), *elav-Gal4* (*3E1*, BDSC #8760), and *Sirt1*^*2A-7-11*^ (BDSC #8838).

### RNA Measurement

RNA was extracted from male heads, DNase treated, and reverse-transcribed using MultiScribe™ (Applied Biosystems). Quantitative PCR reactions were done using the SYBR Green method and custom designed primers on a StepOnePlus machine (Applied Biosystems). C_t_ values were normalized to *RpL32*, expression was calculated using the ΔΔC_t_ method, and the mean of multiple independent biological replicates was calculated. Oligonucleotide primers used in this study were

*Cdc7:* AATGGAGCTGCAGTCATGG (F),

GGATTCGTGTGAGGAGATCATT (R); *CG14186:*

GGCCAGCTAATCTCCAAGTT (F), GTTGTAGATCTCCTCGCCATC

(R); *CG17778:* GCTGCGCTGACTTACTACTTAC (F),

TGCATTGGCCACCGATTT (R); *Hr38:* GAGTGGCTCAACGACATCAT

(F), CGTTCTGTGATCAGGGTTAGG (R); *Jra*:

GTTCCCACCCACTGATTGA (F), GCTTGTTCTTGGCACTCTTG (R);

*Kayak:* CCGATACTTCAAGTGCCCATAC (F),

CCAGGACATTGGAGAAGTTGTT (R); *Sirt1:*

GACTGCCGGATGAGTACC (F), ACGATCAGTAGATCGCAC (R);

*Stripe:* CCGAGTATGCCGCTCAATTA (F),

GGCGTATGGTGGTGATAAGG (R).

### Whole Mount Immunohistochemistry

Brains were dissected in PBS and 0.05% Triton-X 100 (0.05% PBT), fixed (2% paraformaldehyde in 0.05% PBT) overnight at 4°C or 1 hr at room temperature. Brains were washed 5x 10 min in 0.1% PBT, blocked 1 hr in 0.1% PBT with 0.5% w/v BSA and 5% normal goat serum and then incubated with primary antibodies overnight at 4°C. Brains were washed, blocked, and incubated with secondary antibodies overnight at 4°C, followed by further washes and then mounted on glass slides with Vectashield (Vector Laboratories). Antibodies used were rabbit anti-GFP (1:1000, Invitrogen A6455), mouse anti-Elav (1:50, Developmental Studies Hybridoma Bank 9F8A9), goat antirabbit Alexa 488, and goat anti-mouse Alexa 594 (1:350, Cell Signaling Technologies). Mushroom body kenyon cell nuclei were counted by drawing a 50 μm arc at the border between the mushroom body calyx and nuclei at the location with the greatest number of GFP-positive nuclei, and counting positive nuclei within 25 μm of the border.

### Ethanol Behaviors

Ethanol sensitivity and tolerance were measured as previously described (Engel *et al.* 2016). Briefly, groups of twenty genetically identical flies (n=1) were exposed to 55% ethanol vapor or 100% humidified air, and the number of flies that lost the righting reflex were counted at 6 min intervals. The time to 50% sedation (ST50) was calculated for each group, and the experiment was repeated across different days and from different parental crosses. Flies were allowed to rest for 3.5 hr and then re-exposed to an identical concentration of ethanol vapor, and tolerance was calculated as the difference in ST50 between the two exposures. The capillary feeding assay (CAFE) was used to determine ethanol preference, as previously described (Devineni & Heberlein 2009). Groups of eight adult males were collected 3-4 days after eclosion and allowed to recover from CO_2_ for one day. They were pre-exposed to either 55% ethanol vapor/air mixture or 100% humidified air alone for 20 min. After 16 hr recovery, flies were placed into the CAFE chamber, which consists of empty vials with capillary tubes containing liquid food with or without 15% ethanol, embedded in the vial plug. The preference index was the volume of food consumed from the ethanol capillaries minus that consumed from the no-ethanol capillaries over the total volume consumed, corrected for evaporation by measuring the volume lost in vials with no flies. Bitter taste avoidance was measured by presenting flies with a choice of 1.25% agarose containing either 50 mM sucrose (S) or 100 mM sucrose and 1 mM quinine (SQ). Groups of approximately 20 male flies were food deprived on water for 14 h, placed in a 40×90×10 mm clear acrylic arena, and 150 uL S and SQ dots were then placed in apposition at the center of the arena. The number of flies on each dot was counted at 30 min. Avoidance was calculated as (SQ − S)/(SQ + S) such that complete avoidance of bitter gives a value of −1.

### Ethanol Absorption and Metabolism

Flies were frozen in liquid nitrogen and homogenized in 50 mM Tris-HCl, pH 7.5. Ethanol concentrations were measured in fly homogenates using the Ethanol Assay Kit from Diagnostic Chemicals Ltd. (catalog #229-29). To calculate the ethanol concentration in flies, the volume of one fly was estimated to be 1 μl.

### Statistics

GraphPad Prism 7.0c was used for unpaired *t* test, one sample *t* test, one-way ANOVA with Tukey’s *post hoc* test for normally distributed data, and Kruskal-Wallis test with Dunn’s *post hoc* test for non-parametric data. Significance indicators on the figures indicate the results of *t* tests or *post hoc* tests for significant effects by ANOVA. Error bars represent the SEM.

## Results

### Hr38 is induced by acute ethanol exposure

We surveyed a subset of immediate early genes that are broadly induced by neuronal activity to ask if drug naïve Drosophila respond to ethanol transcriptionally through similar pathways (Guan *et al.* 2005; Fujita *et al.* 2013; Chen *et al.* 2016). Of these genes, the Nr4a nuclear hormone receptor homolog Hr38 was the only immediate early transcription factors whose expression was induced to statistical significance (Figure 1a). The Jun-related antigen *Jra* gene showed a trend towards induction. Retrospective analysis of a gene expression time course following acute ethanol exposure revealed that *Hr38* levels peaked 60 min after ethanol exposure termination and then decreased to baseline within three hours, kinetics that are typical for immediate early response genes (Figure 1b). Thus, ethanol induces immediate early genes in a pattern that partially overlaps that of neuronal activation.

**Figure 1.**
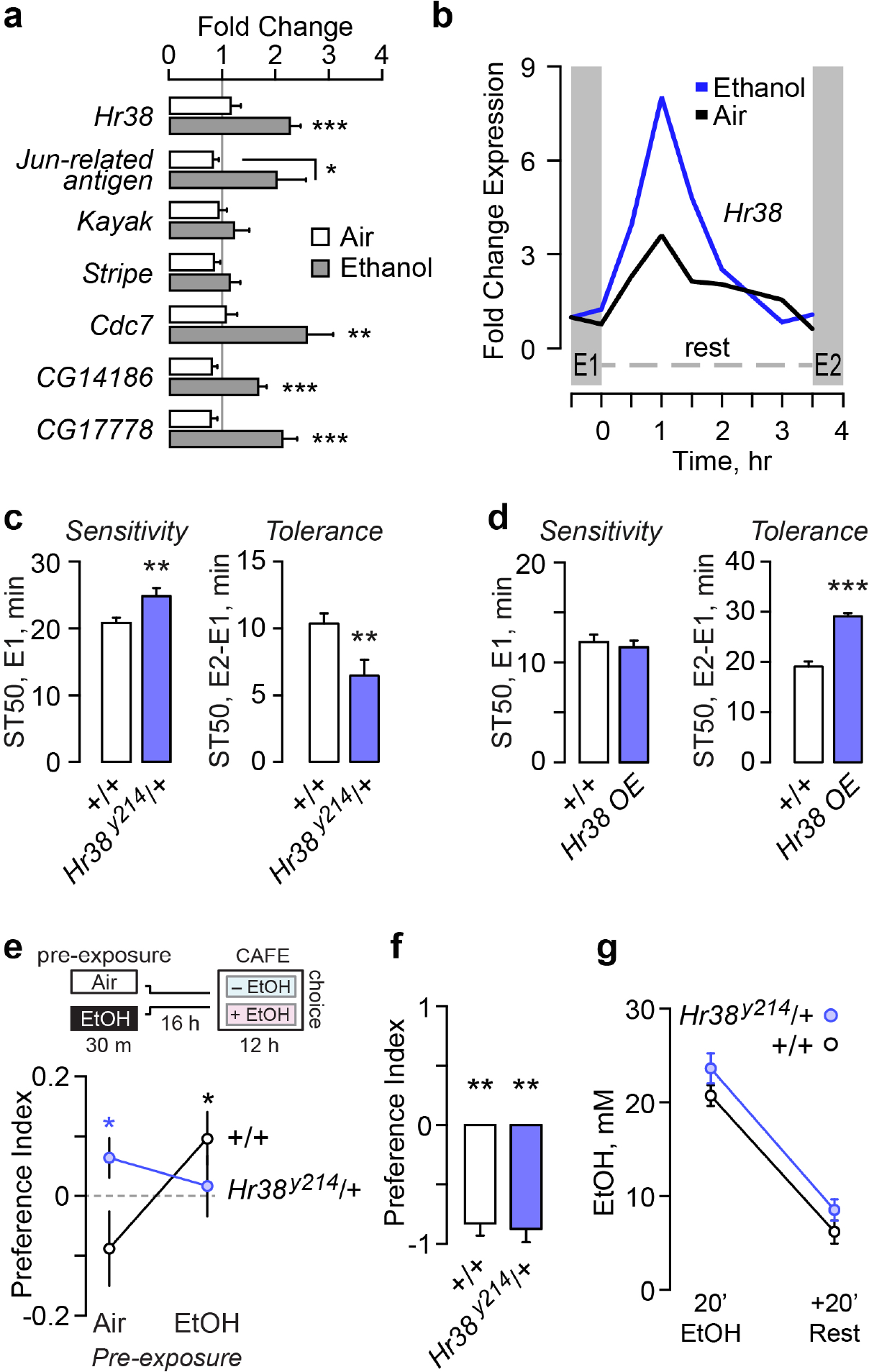
*Hr38* is an ethanol immediate early response gene that bidirectionally regulates the development of ethanol tolerance. **a**. Steady state transcript levels of genes induced by neuronal activity, 60 min after termination of 30 min ethanol or mock air exposure, presented as fold change vs. no treatment (grey line) in the wild-type control strain. One way ANOVA/Tukeys per gene, n=5 biological replicates/condition. **b**. Time course of *Hr38* expression following air (black) or ethanol (blue). Data extracted from a published microarray experiment (Kong *et al.* 2010). **c**. Time to 50% sedation (ST50) for *Hr38* null mutant heterozygotes vs. wild-type controls (+/+) exposed to ethanol once (E1) for sensitivity or twice (E2-E1) for tolerance. t-test, n=30 groups. **d**. Ethanol sensitivity and tolerance in flies with three copies of the *Hr38* genomic region (*HR38.GFP*). t-test, n=12 groups. **e**. Ethanol preference in *Hr38* null heterozygotes. Flies are pre-exposed to either air or ethanol and after 16 hr placed into the 2 choice CAFÉ assay. A positive index indicates preference for ethanol intake. One-tailed t-test compared to zero preference, n=20 groups. **f**. *Hr38* null heterozygotes show avoidance of a bitter but sweeter food source in a two-choice seeking assay. One sample t-test compared to zero preference, n=5 groups. **g**. Ethanol accumulation immediately following a 20 min exposure, and after a 20 min rest, allowing ethanol to be metabolized. t-test, n=8 groups. *P<0.05, **P<0.01, ***P<0.001.

### Hr38 promotes ethanol tolerance and ethanol preference

Induction of the transcription factor Hr38 suggested that it may regulate gene expression in the nervous system to promote ethanol behavioral plasticity. To ask if Hr38 functions in ethanol behaviors, we tested flies underexpressing or overexpressing the gene for ethanol sensitivity, ethanol rapid tolerance, and ethanol preference. Ethanol sensitivity was measured as the time to 50% sedation for groups of genetically identical flies. Ethanol tolerance was measured by giving these flies a second, identical ethanol exposure 3.5 hours after the first exposure, and calculating the difference in sedation time between exposures: flies acquire resistance to the sedative effects of ethanol (Scholz *et al.* 2000). *Hr38*^*y214*^ null mutants are pupal lethal, so we tested heterozygotes with 50% normal *Hr38* levels. *Hr38* heterozygotes showed decreased ethanol sensitivity and decreased ethanol tolerance (Figure 1c). A BAC insertion of the *Hr38* genomic region, when heterozygous in wild-type flies, increased *Hr38* genomic copy number from two to three and expression 1.78 fold (p=0.0154 two-tailed t-test, n=6 biological replicates). *Hr38* overexpression did not affect ethanol sensitivity, but it markedly increased ethanol tolerance (Figure 1d). This suggests that Hr38 levels are critical for setting the magnitude of ethanol tolerance.

Drosophila develop a preference for ethanol intake (Devineni *et al.* 2011). Ethanol preference was measured in the two choice CAFÉ assay, where flies can drink from capillaries containing sucrose and yeast either with or without 15% ethanol (Ja *et al.* 2007; Devineni & Heberlein 2009). Ethanol preference in wild-type was induced by pre-exposure to ethanol vapor (Figure 1e) (Peru Y Colón de Portugal *et al.* 2014). In contrast, *Hr38* loss-of-function mutants showed precocious ethanol preference, and this preference dissipated with ethanol pre-exposure (Figure 1e). Bitter taste avoidance was unaffected in the *Hr38* mutants, suggesting that their ethanol taste reactivity is intact (Figure 1f). Moreover, ethanol absorption and metabolism were unaffected in *Hr38* mutants (Figure 1g). Taken together, these results suggest that the levels of *Hr38* expression are important for two forms of ethanol behavioral plasticity, tolerance and preference.

**Figure 2.**
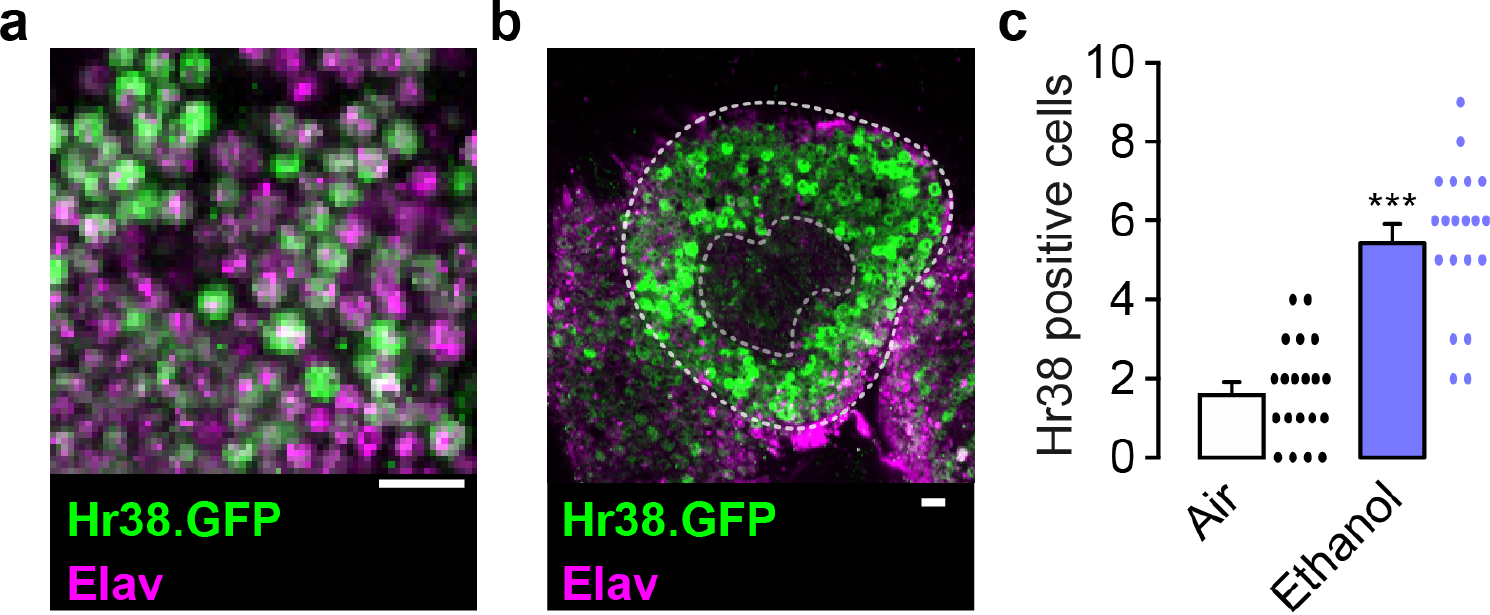
Hr38 protein expression in the adult brain is induced by ethanol in the mushroom bodies. **a**. Cortex region of the adult fly brain expressing the Hr38.GFP fusion protein (green), showing co-localization with the pan-neuronal Elav nuclear protein (magenta). **b**. High levels of Hr38.GFP expression in the Kenyon cell neurons (between dashed enclosures) of the mushroom body that surround the mushroom body calyx (inner dashed area) in the brain of an ethanol exposed fly. **c**. Number of Hr38.GFP-positive Kenyon cell nuclei 1.5 hr after termination of ethanol or mock air exposure. ***P<0.0001, t-test, n=20 brains/treatment. Scale bars: 10 μm.

### Hr38 is expressed in neurons where it is induced by ethanol

Hr38 on the genomic BAC is C-terminal tagged with GFP (Hr38.GFP), which allowed us to determine its expression pattern in the fly brain (Figure 2). Hr38.GFP was localized to the cell nucleus but was also present in the cytoplasm in some brain regions, most notably in the axons of the mushroom body lobes (Figure 2a, Supplemental Movie 1). Co-labeling indicated that Hr38-positive cells were neurons (Figure 2a). Hr38.GFP expression was found sporadically throughout the adult brain, and the mushroom body kenyon cell nuclei were prominently labeled (Figure 2b). To ask if ethanol exposure recruited additional neurons to express Hr38, we counted GFP-positive Kenyon cell nuclei. The number of Hr38-labeled nuclei increased about 2-fold in the mushroom body 1.5 hr after termination of ethanol treatment (Figure 2c).

### Mef2 acts upstream of Hr38 for ethanol tolerance in the mushroom bodies

Mammalian homologs of Hr38 are transcriptionally induced by the Mef2 transcription factor (Flavell *et al.* 2006; Shalizi *et al.* 2006). We asked if ethanol upregulates *Hr38* through Mef2 to promote tolerance development. Mef2 consensus DNA binding sites (CTAWWWWTAG) are overrepresented in the *Hr38* genomic region, two of which are located less than 2 kb upstream of the *Hr38* transcription start site (Figure 3a) (Andrés *et al.* 1995; Sivachenko *et al.* 2013). Two of the Mef2 consensus sites bind Mef2 in chromatin immunoprecipitation from fly heads, and they are also conserved in *Drosophila simulans* (Sivachenko *et al.* 2013). Consensus sites also exist in the *Apis melifera Hr38* enhancer region (not shown). We used a transgenic dominant negative Mef2, Mef2.EnR, to inhibit transcriptional activity at Mef2 enhancers in neurons (Blanchard *et al.* 2010). In Mef2.EnR, the Mef2 activation domain is replaced with the Engrailed repressor domain. Expression of Mef2.EnR in all neurons with *elav-Gal4* reduced *Hr38* expression (Figure 3b). Further, *Hr38* induction by ethanol was lost (Figure 3c). These data suggest that Mef2 is an immediate early activator of *Hr38* gene transcription, and that acute ethanol exposure acts at or upstream of Mef2 to change gene expression in neurons.

These findings predicted that decreased Mef2 activity in neurons would decrease ethanol tolerance. We used two methods to reduce Mef2, RNAi (*Mef2.IR*) and Mef2.EnR. Mef2 RNAi in all neurons, like *Hr38* loss-of-function mutants, decreased both ethanol sensitivity and ethanol tolerance (Figure 3d). A second *Mef2* RNAi appeared to be weaker, specifically reducing ethanol tolerance (not shown). Dominant negative Mef2 in all neurons decreased ethanol tolerance, but had no effect on ethanol sensitivity (Figure 3e).

The α/β lobe neurons of the mushroom bodies promote ethanol tolerance, where Mef2 and Hr38 expression are enriched (Schulz *et al.* 1996; Engel *et al.* 2016). Expression of dominant negative Mef2 in these neurons using the *17d-Gal4* mushroom body driver reduced ethanol tolerance (Figure 3f). Co-expression of the Gal4 repressor Gal80 specifically in the mushroom bodies blocked the effect of Mef2.EnR on ethanol tolerance, indicating that Mef2 promotes tolerance in the mushroom bodies (Figure 3g).

**Figure 3.**
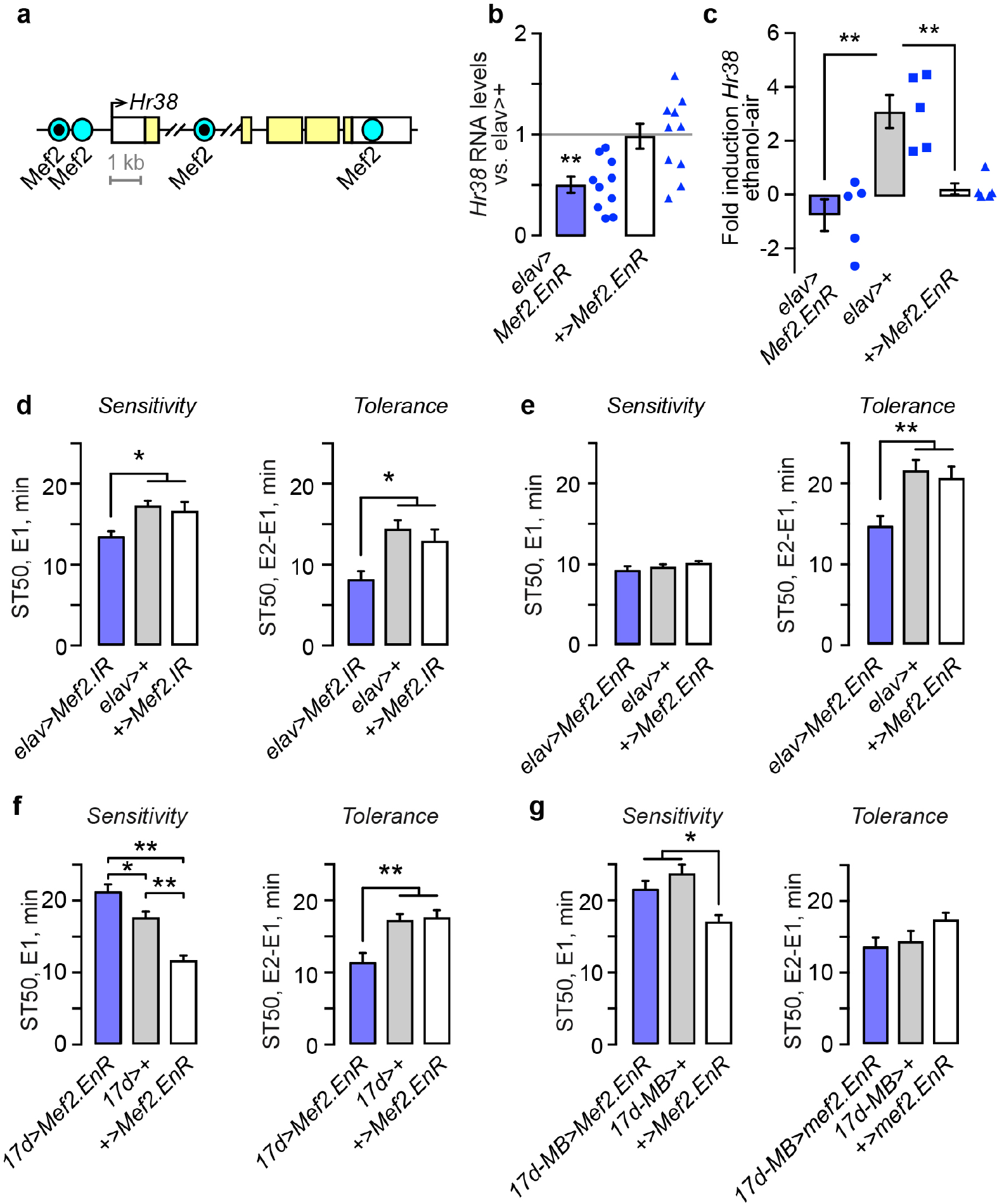
Mef2 promotes ethanol induction of *Hr38*, and ethanol tolerance in the mushroom bodies. **a**. Diagram of the *Hr38* genomic locus, indicating the positions of Mef2 consensus binding sites. Consensus sites conserved in other Drosophila species are indicated by a black dot. **b**. *Hr38* transcript levels in flies expressing dominant negative Mef2.EnR panneuronally, normalized to elav-3E1>+ (grey line). Kruskal-Wallis/Dunn’s, n=10 biological replicates. **c**. Ethanol induction of *Hr38* 60 min after ethanol treatment is blocked by Mef2.EnR. Expressed as the difference between ethanol and mock air exposure per biological replicate. No difference was detected between genotypes for the mock air exposure. One-way ANOVA/Tukey’s, n=5. **d**. Ethanol sensitivity and tolerance with pan-neuronal expression (*elav-c155*) of Mef2 RNAi (*Mef2.IR*). One way ANOVA/Tukey’s, n=30 groups. **e**. Ethanol sensitivity and tolerance with pan-neuronal expression of *Mef2.EnR*. One way ANOVA/Tukey’s, n=14 groups. **f**. Ethanol sensitivity and tolerance with expression of Mef2.EnR restricted to the mushroom body α/β lobes (*17d-Gal4*). One way ANOVA/Tukey’s, n=15-16 groups. *P<0.05, **P<0.01. **g**. Ethanol sensitivity and tolerance in 17d>*Mef2.Enr* in the presence of the mushroom body-specific *MB-Gal80*. One way ANOVA/Tukey’s, n=10-12 groups. *P<0.05, **P<0.01.

### Sirt1 terminates ethanol-induction of Hr38

Sirt1 (also known as Sir2) is a histone/protein deacetylase that regulates responses to drugs of abuse in Drosophila and mammals (Renthal *et al.* 2009; Engel *et al.* 2016). In Drosophila, mushroom body α/β lobe core neuron promotion of ethanol tolerance, preference, and reward requires *Sirt1*. Further, *Sirt1* broadly allows gene expression regulation by acute ethanol exposure. We therefore asked if absence of *Sirt1* affected ethanol induction of *Hr38*. *Hr38* was induced normally at 1 hr after acute ethanol exposure in *Sirt1* null mutants (Figure 4a). Because deacetylation of histones by Sirt1 is generally thought to compact chromatin and decrease gene expression, we also assessed *Hr38* expression 3 hr after ethanol termination, when *Hr38* levels have returned to pre-exposure levels. *Hr38* expression was markedly higher in *Sirt1* null mutants at 3 hr (Figure 4a). Thus, *Sirt1* is required for termination of *Hr38* gene expression induction by acute ethanol exposure. Finally, we detected no change in *Sirt1* levels with neuronally expressed dominant negative Mef2, indicating that *Mef2* and *Sirt1* act independently to regulate *Hr38* expression (Figure 4b).

**Figure 4.**
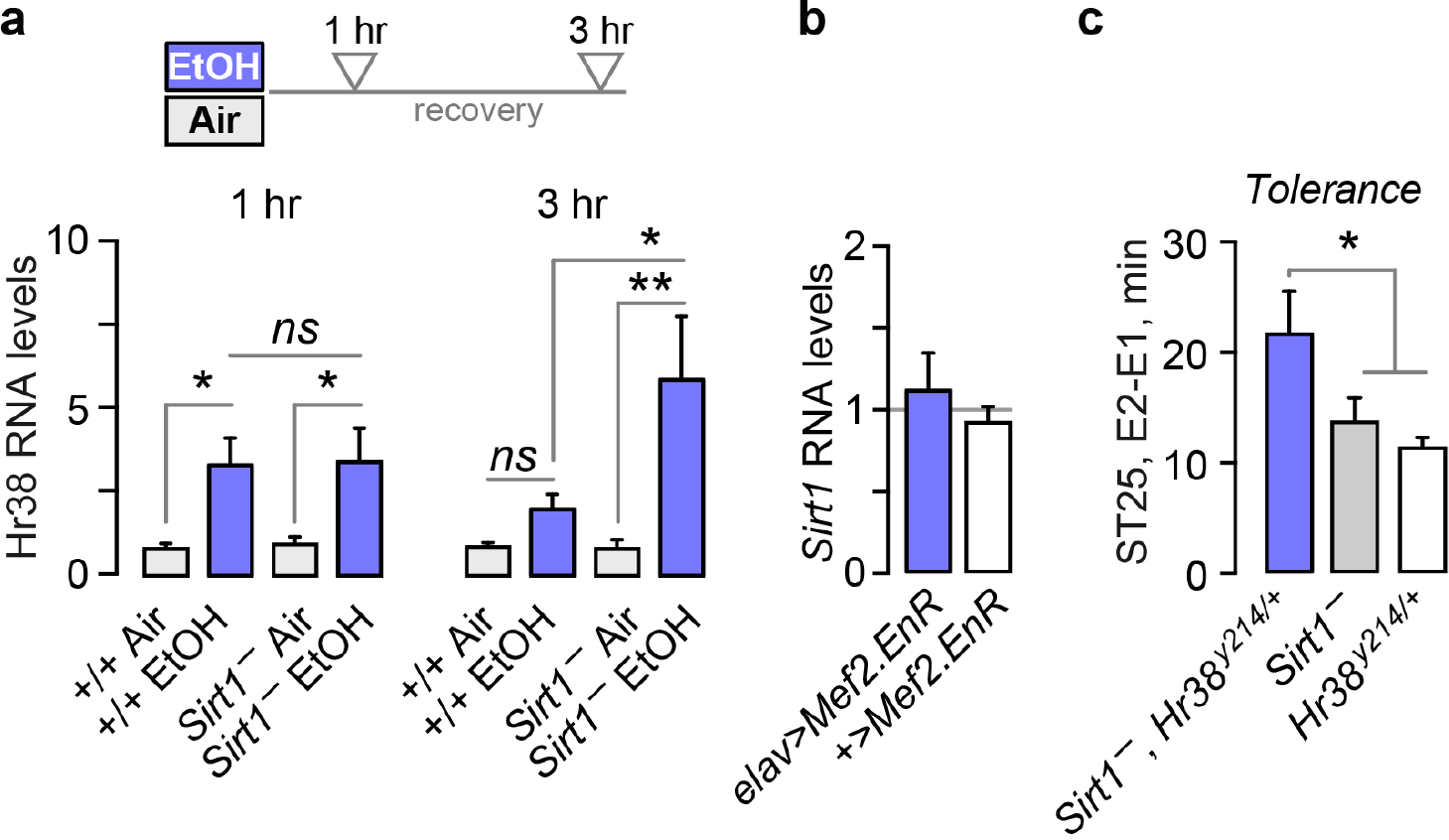
*Hr38* interactions with *Sirt1*. **a**. *Hr38* transcript expression levels in wild-type controls (+/+) and *Sirt1* null mutants 1 hr and 3 hr after treatment. One way ANOVA/Tukey’s, n=14. **b**. *Sirt1* transcript levels in flies expressing dominant negative Mef2 in all neurons (*elav-3E1*). Kruskal-Wallis/Dunn’s, n=4-5. **c**. Ethanol tolerance in flies null for *Sirt1* and heterozygous for *Hr38*. Because of the high tolerance in the double mutants, the time to 25% sedation (ST25) was measured. One way ANOVA/Tukey’s, n=7-13. *P<0.05, **P<0.01.

Induced *Hr38* may need to be terminated rapidly in order for behavioral plasticity to proceed. We performed a test of this by making double mutants with *Sirt1* and *Hr38*, predicting that genetically decreasing *Hr38* expression by half may moderate Hr38 induction in Sirt1 mutants and allow ethanol tolerance to develop. The double mutants showed increased tolerance, compared to either mutant alone (Figure 4c). This data suggests that termination of Hr38 expression by Sirt1 is important for the development of ethanol tolerance.

## Discussion

How ethanol changes gene expression in the nervous system is a key to understanding how ethanol causes maladaptive changes in brain function. Here we show that ethanol exposure in drug naïve Drosophila causes a Mef2-dependent increase in Hr38 expression, that it is terminated by Sirt1, and that Hr38 controls the extent of ethanol tolerance development. These data suggest that ethanol acts upstream of or directly on the Mef2 transcription factor to cause gene expression changes in the nervous system. Our studies also reveal a requirement for temporally precise termination of the immediate early transcriptional response. Genes regulated by the Hr38 transcription factor are candidate effectors for ethanol-induced behavioral plasticity.

In both flies and mammals, acute ethanol exposure causes changes in both gene expression and chromatin structure in the brain. In Drosophila, transcriptomic experiments that used varying ethanol exposures and recovery times on whole head samples discovered over 200 common ethanol responsive genes (Morozova *et al.* 2007; Urizar *et al.* 2007; Kong *et al.* 2010). Similarly, ethanol exposure increases histones H3 and H4 acetylation marks that generally indicate the opening of chromatin (Ghezzi *et al.* 2013; Engel *et al.* 2016). Like gene expression, histone H4 acetylation was modified at a large number of genomic loci 6 and 24 hr after ethanol exposure (Park *et al.* 2017). Further, the increase in histone acetylation can be quite rapid, starting during inebriation (Engel *et al.* 2016). Therefore, acute ethanol induces genomic changes rapidly, broadly, and lastingly. Hr38 and the other ethanol-induced immediate early response genes we identified are candidates for controlling major aspects of ethanol neuroadaptation. In particular, Hr38 as a transcription factor may control the expression of downstream effector genes. The reciprocal effects on ethanol tolerance of lowering and raising Hr38 levels, plus the consequences of prolonging Hr38 expression, all argue that Hr38 is a key regulator of the genomic program for ethanol neuroadaptation.

Hr38, as the sole homolog of the mammalian Nr4a1/2/3 gene family, may carry out some of the functions ascribed to distinct mammalian family members. Mammalian Nr4a transcription factors are induced by neuronal activity, stress, and drugs of abuse (Campos-Melo *et al.* 2013). In particular, Nr4a1, also known as Nurr77 and NGFIB, is upregulated in by cocaine and morphine in brain regions implicated in addiction, and deletion of Nr4a2, also known as Nurr1, decreases ethanol preference (Werme *et al.* 2000, 2003; Chandra *et al.* 2015). Nr4a1 and Nr4a2 are also implicated in forms of long-term memory (McNulty *et al.* 2012). Nr4a1 induced by neuronal activity regulates the density and distribution of dendritic spines, suggesting a possible link between acute ethanol activation of Hr38 and changes in the functional connectivity of the brain (Chen *et al.* 2014).

Hr38 in Drosophila controls cuticle development and glycogen storage, and its expression is regulated by neuronal activity, social cues, light, and extreme drops in temperature (Kozlova *et al.* 2009; Ruaud *et al.* 2011; Fujita *et al.* 2013; Adewoye *et al.* 2015; von Heckel *et al.* 2016). Hr38’s role in ethanol responses is likely distinct from most of these roles, since manipulating its levels specifically in the nervous system affects the development of tolerance, and since we detected no change in ethanol absorption as would be expected for altered cuticle integrity (Singh & Heberlein 2000). Further, we showed that blocking Mef2 activity in the mushroom bodies, where Hr38 expression is increased by ethanol, decreases ethanol tolerance. Mushroom body induction of Hr38 also occurs in males when presented with females, suggesting a possible molecular and anatomical link between ethanol and sexual behaviors (Fujita *et al.* 2013).

How is Hr38 expression controlled? We show that Mef2 sets Hr38 levels under steady state conditions and promotes Hr38 induction by ethanol. In mammals, Mef2A and Mef2D both increase Nr4a1 expression to regulate synapse number and dendrite differentiation (Flavell *et al.* 2006; Shalizi *et al.* 2006). Further, Mef2C and NR4a1 are coordinately increased in the striatum by cocaine, and Mef2 promotes cocaine sensitization in the nucleus accumbens, a brain region critical for drug reward (Pulipparacharuvil *et al.* 2008; Dietrich *et al.* 2012). Mef2 expression and activity can be regulated by many different signaling pathways, including those associated with neural activity like intracellular calcium levels, and also those that are known to be regulated by ethanol in mammals (Flavell *et al.* 2006; Hawk & Abel 2011; Ron & Barak 2016). The concomitant upregulation by ethanol of numerous other immediate early genes suggests that acute ethanol may act in part through neuronal activity pathways (Chen *et al.* 2016).

Sirt1 is broadly required for ethanol induction of gene expression in flies, and we show here that it is also required to terminate *Hr38* expression after *Hr38* is induced by ethanol (Engel *et al.* 2016). *Hr38* induction by ethanol, however, appears to be independent of Sirt1. This may be due to the distinct chromatin structure at immediate early gene loci (Chen *et al.* 2017). Moreover, the loss of Sirt1 deacetylase activity in Sirt1 mutants may directly result in prolonged acetylation of the Hr38 locus and its continued transcriptional activation. In wild-type flies, ethanol causes a transient decrease in Sirt1 gene expression and protein levels that temporally coincides with termination of Hr38 gene expression. This suggests that a more complex set of molecular events involving Sirt1 terminates *Hr38* expression. This may involve other chromatin modifying enzymes, other genes whose expression is regulated by Sirt1, or one of the non-histone targets of Sirt1 (Gomes *et al.* 2016; Yang *et al.* 2016).

## Acknowledgments

We thank Harpreet Randhawa for technical assistance, the Wolf and Michael Cleary lab members and Ramendra Saha for discussions, and the Bloomington Drosophila Stock Center for strains. This work was supported by grants to F.W.W. from the NIH/NIAAA R03AA023262 and R21AA025560. The authors state no conflict of interest.

